# Tracking crystallophore nucleating properties: setting-up a database for statistical analysis

**DOI:** 10.1101/2020.04.23.057596

**Authors:** Tao Jiang, Amandine Roux, Sylvain Engilberge, Zaynab Alsalman, Sebastiano Di Pietro, Bruno Franzetti, François Riobé, Olivier Maury, Eric Girard

## Abstract

In this article, the principle of a database aimed at facilitating the understanding of the unique protein nucleating properties of the Crystallophore is presented. A first analysis allows us to compare the efficiency of Tb-Xo4 with the new Lu-Xo4 variant, featuring improved phasing properties. Then, the concept of *subset-of-interest* is introduced to reveal potential antagonistic/synergistic effects between Tb-Xo4 and physico-chemical parameters of the crystallisation kits such as pH. The overall approach may be of interest for any studies working on solutions dedicated to improve the nucleating step in protein crystallization.

**TOC Graphic:** 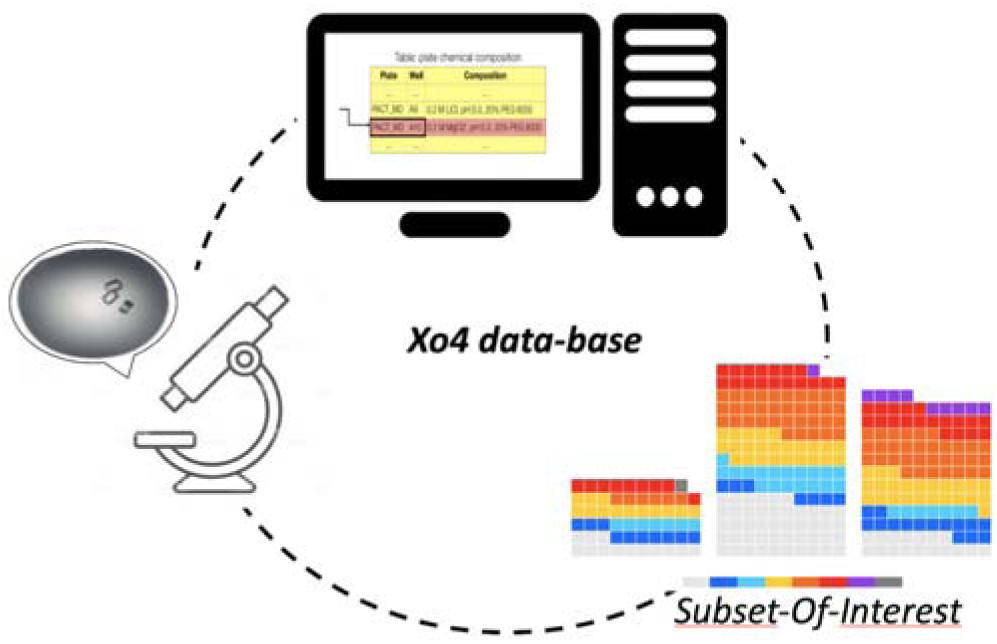

**Synopsis:** A database and associated representation tools are highlighted to understand nucleating properties of the crystallophore.

## INTRODUCTION

The success of protein structure determination by X-ray crystallography obviously relies on the availability of well-diffracting crystal that remains precisely the major hurdle in this multi-steps process. Automation associated with the use of nano-volume dispensing robot has tentatively increased the efficiency of the process. Depending on protein sample availability, hundred to thousand crystallization conditions can be evaluated thanks to the various commercial kits available on the market, without guarantee of success in getting a suitable condition. Different approaches facilitating the nucleation step have been proposed: (i) when an initial crystallization condition provides even poor quality crystals or related like urchins, homogenous nucleation can be envisaged through seeding of crystallization drops with fragments of crystal.^1-4^ (ii) Secondly, heterologous nucleation relies on the introduction of materials within the drop at the start of the crystallization process. Various materials have been evaluated with more or less success.^5-11^ (iii) Finally, the recent development of molecular glues (calixarene, polyoxometalate or lanthanide complexes) able to improve or mediate the protein-protein contacts is nowadays the most straightforward strategy to improve the nucleation step.^12-17^ Among these molecular glues we introduced, in 2017, the crystallophore.^18^

The crystallophore, Xo4, is a family of cationic lanthanide complexes presenting both nucleating and phasing properties in an all-in-one tool which is capable of overcoming the two main bottlenecks of macromolecular crystallography.^18^ The story so far of the Xo4 system started with its terbium variant Tb-Xo4. Its unique properties were firstly highlighted with a set of 8 proteins and its efficiency was confirmed in solving issues often dealt by crystallographer in their quest to assess the protein structure.^18,19^ Finally, Tb-Xo4 was exploited to determine new protein structures including multi-protein complexes^20-22^ and can be used as a routine nucleating and phasing tool by the community.^23-27^

To understand the origins of the crystallophore nucleating and phasing properties with the aim to further improve them, we analyzed the binding of Tb-Xo4 at the surface of proteins by coupling crystal structure analysis, obtained on 4 different proteins, with evaluation of the interaction energies determined by density functional theory calculations.^28^ This detailed analysis of the interaction sites on the proteins surfaces clearly demonstrated the great versatility of Tb- Xo4 binding through various supramolecular interactions via anionic, cationic or hydrophobic amino-acid residues. Such versatility may explain the unprecedented properties of this compound. However, this variability in the interaction mode from one example to another prevents any determination of an optimal amino-acid environment required for an efficient interaction of Tb-Xo4 that could enhance the nucleation process. Furthermore, the study also revealed the non-negligible role of the particular composition of the crystallization solution or protein buffer in the global interaction networks. We observed the direct involvement of components, such as Ca^2+^ cation or glycerol molecule, in the interaction network. Sulfate or iodide anions may also directly interact with crystallophore turning its overall charge (mono-cationic) into a neutral or even an anionic one thus modulating the interaction energies balance.^28^ Thus, it appears clearly that the crystallization solution components may directly participate to the nucleation process making its study much more complicated due to the huge number of independent parameters.

In order to understand how crystallization kits formulation and corresponding physico-chemical conditions may influence the nucleating effect of the crystallophore, we set-up a database to efficiently analyze the influence of crystallization media components. In the present paper, we present the structure of this database through a first analysis of 6 crystallization trials that were performed in the course of our previous work.^18^ This analysis will benefit from the comparison with the crystallization screening in native conditions that was run in absence of the crystallophore. These 6 proteins (hen egg white lysozyme (HEWL), *Tritirachium album* proteinaseK, *Thaumatococcus daniellii* thaumatin, protease one from *P. horokoshii*, Glyoxylate Hydroxypyruvate Reductase from *Pyrococcus furiosus* (GRHPR) and bacteriophage T5 distal tail protein (pb9)) were evaluated with the same commercial crystallization kits.^18^

Lanthanide complexes chemistry also offers the possibility to explore several variants of the crystallophore obtained by chemical modifications of the Xo4 ligand, possibly altering interactions properties of the generated variant and thus nucleating properties. In the present work, we introduce a variant of the crystallophore, where the terbium (III) ion was substituted with lutetium (III) (Scheme 1). Indeed, based on a previous study,^29^ the lutetium crystallophore, Lu-Xo4, was synthetized in order to provide a phasing agent more convenient to use. The lutetium L_III_ absorption edge, at 1.34 Å, is more easily accessible to beamlines dedicated to protein crystallography at synchrotron facilities (for comparison L_III_ absorption edge of terbium is located at 1.65 Å). Indeed, when data resolution is limited by the sample environment, in particular the detector size and/or the crystal-to-detector distance, it may be advantageous to work at a shorter wavelength. Moreover, since anomalous effects are more important at high resolution, this gain facilitates the phasing step, making lutetium the most interesting lanthanide. Lutetium still conserves a large anomalous signal (with an anomalous contribution *f”* of about 10 electrons) at the selenium K absorption edge and can be exploited for non-optimal anomalous-based experiment. It thus became clear to check if Lu-Xo4 retained the same nucleating properties as Tb-Xo4.

Herein, we present the overall structure of the database and different representation modes of the information stored in it and their exploitation to compare, for example, the Tb-Xo4/Lu-Xo4 nucleating properties. In addition, exploiting the database to analyse the nucleating performance of Tb-Xo4 even performed on a limited number of proteins also show a potential influence of the pH as well as a potential synergy between Salt-Grid (Hampton Research) and PACT (Molecular Dimensions) crystallization kits.

**Scheme 1:**
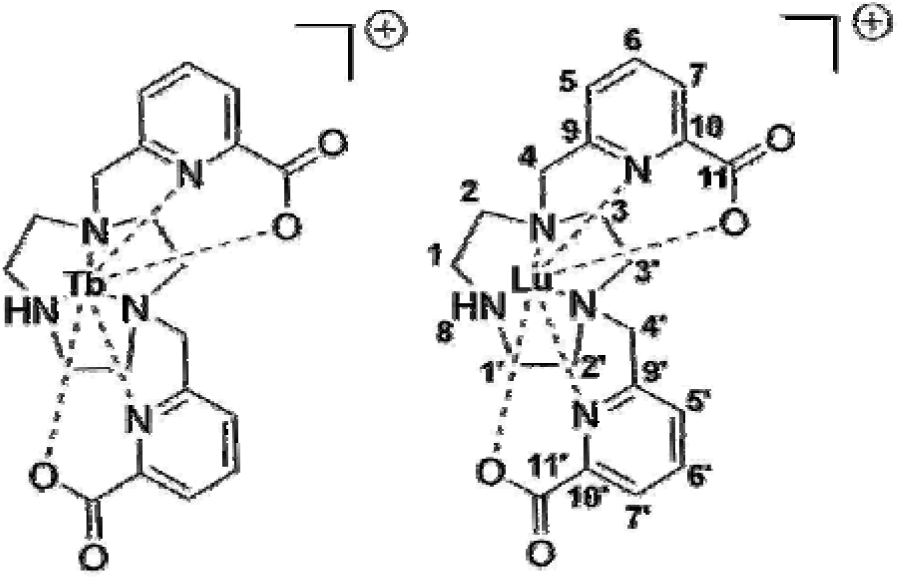
Molecular structure of the crystallophore additives Tb-Xo4 and Lu-Xo4 with atom numbering.

## EXPERIMENTAL SECTION

### Crystallophore synthesis and purification

Tb-Xo4 and Lu-Xo4 were produced using the same improved protocol.^18^ The acid ligand was formed *in situ* from 502 mg of diester (1.17 mmol) in presence of NaOH (2 eq.) in a H_2_O/MeOH mixture (9/1, v/v) stirred overnight at room temperature before the complexation step (HR-MS (ESI) calculated m/z = 400.1979 [M+H]^+^, measured m/z = 400.1975 for M = C_20_H_26_N_5_O_4_).

Then the crude acid ligand formed *in situ* was neutralized to pH = 7 adding HCl (1M) followed by the addition of 481 mg of TbCl_3_.6H_2_O (1.29 mmol, 1.1 eq) to the mixture. The pH was increased until pH = 5.5 using NaOH (1M) and the resulting solution was stirred overnight at room temperature. After evaporation of the solvents, the Tb-Xo4 complex was purified using preparative HPLC separation. The purification was performed using a Nucleodur® (Macherey-Nagel) C18 HTec, 5 μm, Preparative VarioPrep 250 mm column at 5 mL/min. Mobile phase consisted in a gradient of solvent A (0.1% ammonium formate in CH_3_CN) and B (0.1% ammonium formate in H_2_O). Method 1 was used: 5% A during 2 min followed by a 5 to 100 % A gradient in 14 min at 5 mL/min. Then the method carried on during 2 min with 100% A followed by a 100 to 5% A gradient in 8 min. (Retention time = 17 min). A white powder is obtained after freeze-drying with a yield of 78% (505 mg). HR-MS (ESI) calculated m/z = 574.1104 [M+H_2_O+H]^+^, measured m/z = 574.1106 for M = C_20_H_25_N_5_O_5_Tb.

The same protocol was used to synthesize the Lu-Xo4 starting from LuCl_3_.6H_2_O with a yield of 49%. The preparative column used for HPLC purification was an Agilent ZORBAX SB-C18, 5 μm, 100 mm. The purification was done with the same Method 1 as for Tb-Xo4 (Retention time = 11min). HR-MS (ESI) calculated m/z = 590.1258 [M+H_2_O+H]^+^, measured m/z = 590.1239 for M = C_20_H_25_N_5_O_5_Lu.

The NMR analysis (Scheme 1) is: ^1^H-NMR (300 MHz, D_2_O): δ 8.31 (dd, *J*= 7.8 Hz, 1H, H6), 8.25 (dd, *J*= 7.8 Hz, 1H, H6’), 8.17 (d, *J*= 7.7 Hz, 1H, H5), 7.93 (d, *J*= 7.7 Hz, 1H, H5’), 7.86 (dd, *J*= 7.2 Hz, 2H, H7/7’), 4.45 (AB system, δ_A_ = 4.51, δ_B_ = 4.39, J_AB_ = 16 Hz, v_A_ = 1352 Hz, v_B_ = 1315 Hz, 2H, H4), 4.25 (AB system, δ_A_ = 4.29, δ_B_ = 4.20, J_AB_ = 16 Hz, v_A_ = 1287 Hz, v_B_ = 1261 Hz, 2H, H4’), 4.12 (broad s, 1H, NH), 3.74 (t, *J*= 12.9 Hz, 1H, H3’), 3.60 (m, 2H, H1’/H1), 3.41 (m, 1H, H3), 3.10 (t, *J*= 17.0 Hz, 1H, H3’), 3.09 (t, *J*= 17.0 Hz, 1H, H3), 3.05 (m, 1H, H1’), 2.86 (m, 1H, H2’), 2.82 (m, 1H, H1), 2.59 (m, 2H, H2/H2’), 2.30 (t, *J*= 12.3 Hz, 1H, H2).^13^C-NMR (100 MHz, D_2_O): δ 172.4 (C11), 172.1 (C11’), 158.4 (C10), 155.7 (C10’), 149.3 (C9), 148.6 (C9’), 142.8 (C6), 142.3 (C6’), 126.6 (C7), 126.4 (C7’), 124.1 (C5), 124.0 (C5’), 65.7 (C4), 64.3 (C4’), 57.0 (C3), 56.6 (C2), 54.7 (C3’), 54.0 (C1), 49.0 (C2’), 45.8 (C1’).

Both complexes can be purchased from the company Polyvalan (Lyon, France).

### Protein production and crystallization

Purchase, production and purification of the six proteins used in the present study (Table 1) as well as their crystallization using the High-throughput Crystallization facility (HTXlab) at EMBL-Grenoble were previously described.^18^ In brief, each protein was evaluated in the absence and in the presence of 10 mM Tb-Xo4 and crystallization assays were performed in 576 conditions from different commercial crystallization kits. Reported crystallization drop evaluation was performed at 90 days.

**Table 1:** Proteins used in this study and their properties.

| Protein | Uniprot entry | No. of residues | Theoretical pI | No. of Asp/Glu | No. of Lys/Arg |
| --- | --- | --- | --- | --- | --- |
| HEWL | P00698 | 129 | 9.32 | 9 | 17 |
| ProteinaseK <sup>1</sup> | P06873 | 279 | 8.25 | 18 | 20 |
| Thaumatin <sup>2</sup> | P02883 | 207 | 8.46 | 18 | 23 |
| Protease1 <sup>3</sup> | O59413 | 166 | 6.11 | 24 | 22 |
| GRHPR <sup>4</sup> | Q8U3Y2 | 336 | 6.24 | 52 | 50 |
| pb9 <sup>5</sup> | Q6QGE8 | 217 | 6.15 | 23 | 20 |

<sup>1</sup> from *Parengyodontium album*. <sup>2</sup> from *Thaumatococcus daniellii*. <sup>3</sup> from *Pyrococcus horikoshii*. <sup>4</sup> Glyoxylate Hydroxypyruvate Reductase from *Pyrococcus furiosus*. <sup>5</sup> bacteriophage T5 distal tail protein.

For the comparison of the nucleating properties of Lu-Xo4 with Tb-Xo4, crystallization assays with HEWL, ProteinaseK and Thaumatin were performed. The assays were run at the HTXlab similarly to the initial evaluation of Tb-Xo4 nucleating properties.^18^ Drop observation was performed after 15 days.

Crystallization conditions marked as hit correspond to drops containing well-defined crystals.

### Database

The data are managed in a SQLite database. The SQLite3 Python library was used to interface with SQLite database for easy data query and modification. Usage of Python scripts is exploited as the data obtained from the query can be easily analyzed with the assistance of the Numpy library. Various visual representations can be generated by utilizing the Matplotlib library.

## RESULTS AND DISCUSSION

### Database structure

The overall structure of the database is illustrated in Figure 1. The data are stored in two tables. The *samples* table stores the assays performed with the name of the protein and if the assay was performed in the absence or in the presence of the crystallophore, through the protein column which possesses the structure XXX_YYY with XXX is the considered protein and YYY refers to native (absence of crystallophore) or to the presence of crystallophore. The hit tracking results are stored in the *samples* table. The *plate chemical composition* table contains all the detailed information of all well components of the different commercial crystallization kits. As highlighted in Figure 1, both *samples* and *plate chemical composition* tables possess the columns Plate and Well. The two tables can be joined together via these two columns to generate on the fly, thanks to a python script, the *protein crystallization condition* table, from which any analysis of the effects of the chemical compositions on the crystallization process can be performed.

**Figure 1:**
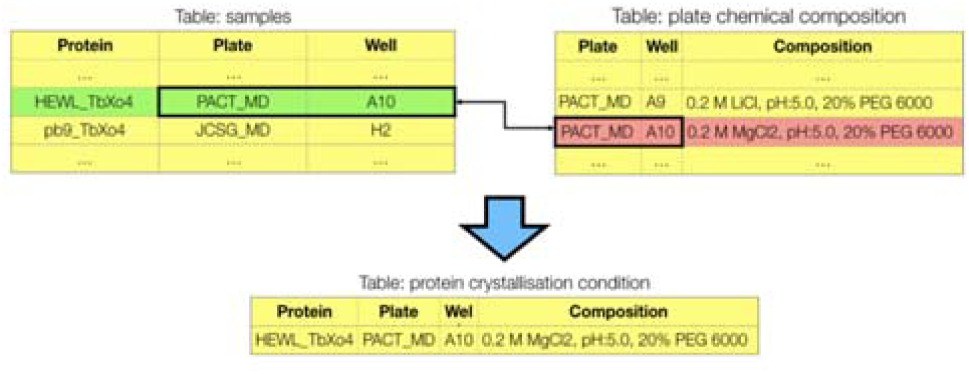
Overall representation of the database structure.

In the framework of studies devoted to understand the crystallization of biological macromolecules, the database offers the unique opportunity to perform rigorous comparative analysis by considering the crystallization of the native protein, *i*.*e*. in the absence of the crystallophore, as a control experiment and to explore the parameters leading to differences induced by Xo4 presence during the crystallization experiment. This exploration will be highly facilitated by the database as experienced by high-throughput crystallization facilities. Indeed, databases are at the heart of such facilities for their daily operation. These databases, as well a the Protein Data Bank, have also been exploited to determine the key factors that affect protein crystallization leading to provide optimized crystallization kits^30-32^ and to promote practical guidelines for initial screening experiments of new crystallization targets.^33-35^

### By-plate representation

In our initial publication,^18^ results of crystallization assays were represented as a histogram providing the number of hits, *i*.*e*. the number of crystallization drops showing crystals, for the native protein (*i*.*e*. in the absence of Tb-Xo4) and for the protein with Tb-Xo4 (10 mM) as well as the number of hits observed for both native and Xo4-containing samples. This representation provides a raw view of the nucleating efficiency of the crystallophore (Figure 2a).

**Figure 2:**
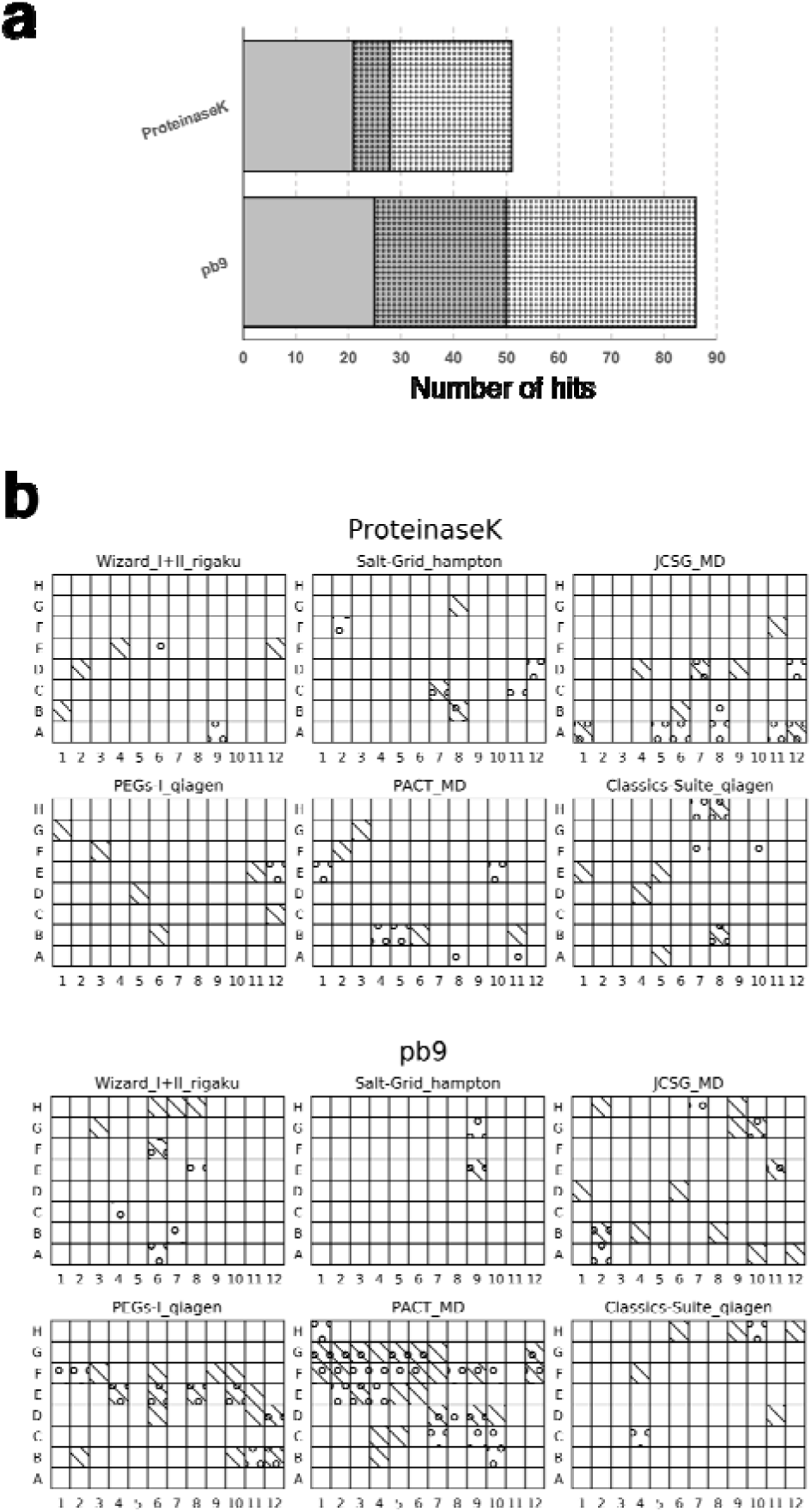
Various representations of the database information for the two proteins, ProteinaseK and pb9. (a) Histogram representation. The number of crystallization hits is depicted in grey for the native protein and with dots for protein supplemented with 10 mM of Tb–Xo4. As a result, the common (shared) conditions are represented in grey with dots. (b) *By-plate* representations where hits without Tb-Xo4 (native) are depicted by a circle while hits obtained in the presence of 10 mM Tb-Xo4 are represented with a backslash.

Thanks to the built database, we explore a *by-plate* representation (Figure 2b). This view allows a side-by-side comparison and also a direct identification of the conditions associated with the crystallization of the native protein alone or with the unique Tb-Xo4 crystallization ones. This view may be of interest for a user looking at tendencies in term of crystallization components in the absence/presence of crystallophore to increase the chance to get exploitable crystals. Indeed, as illustrated in Figure 2b, complementary crystallization conditions can be directly identified and crystallization kits, that might have been neglected based on the native protein crystallization results, now show interest, such as the PEG-Ion or the Classic-Suite kits in the case of ProteinaseK and pb9, respectively.

### Comparison of Lu-Xo4 variant with Tb-Xo4

We exploited the database and the *by-plate* representation to evaluate if Lu-Xo4 variant retained the same nucleating properties as Tb-Xo4. This was determined on three commercial proteins as described in the Experimental section and results.

As expected, most of the hits observed with Tb-Xo4 are recovered with Lu-Xo4 (Figure 3a). These common hits lead to crystals with similar habits (Figure 3b, common hits). However, we also observed crystallization conditions leading to crystals only in the presence of Tb-Xo4 or only in the presence of Lu-Xo4. The number of these unique Tb-Xo4 hits is 18, 7 and 6 for HEWL, ProteinaseK and Thaumatin respectively, while it is 47, 18 and 6 for unique Lu-Xo4 hits. Illustrative drops are displayed in Figure 3b. If some of these unique hits corresponds to crystallization conditions close to those observed for common Tb-Xo4 / Lu-Xo4 hits or may be related to the variability commonly observed in crystallization assays, this does not fully explain the large number observed for Lu-Xo4 in the case of HEWL and ProteinaseK. This may reflect a slightly higher nucleating property of Lu-Xo4. Considering the prevalence of the binding of the central lanthanide ion with pendant carboxylate-containing amino-acids in the Ln-Xo4 interaction modes, this improvement could be induced by the stronger Lewis acidity of the lutetium(III) regards to terbium(III).

**Figure 3:**
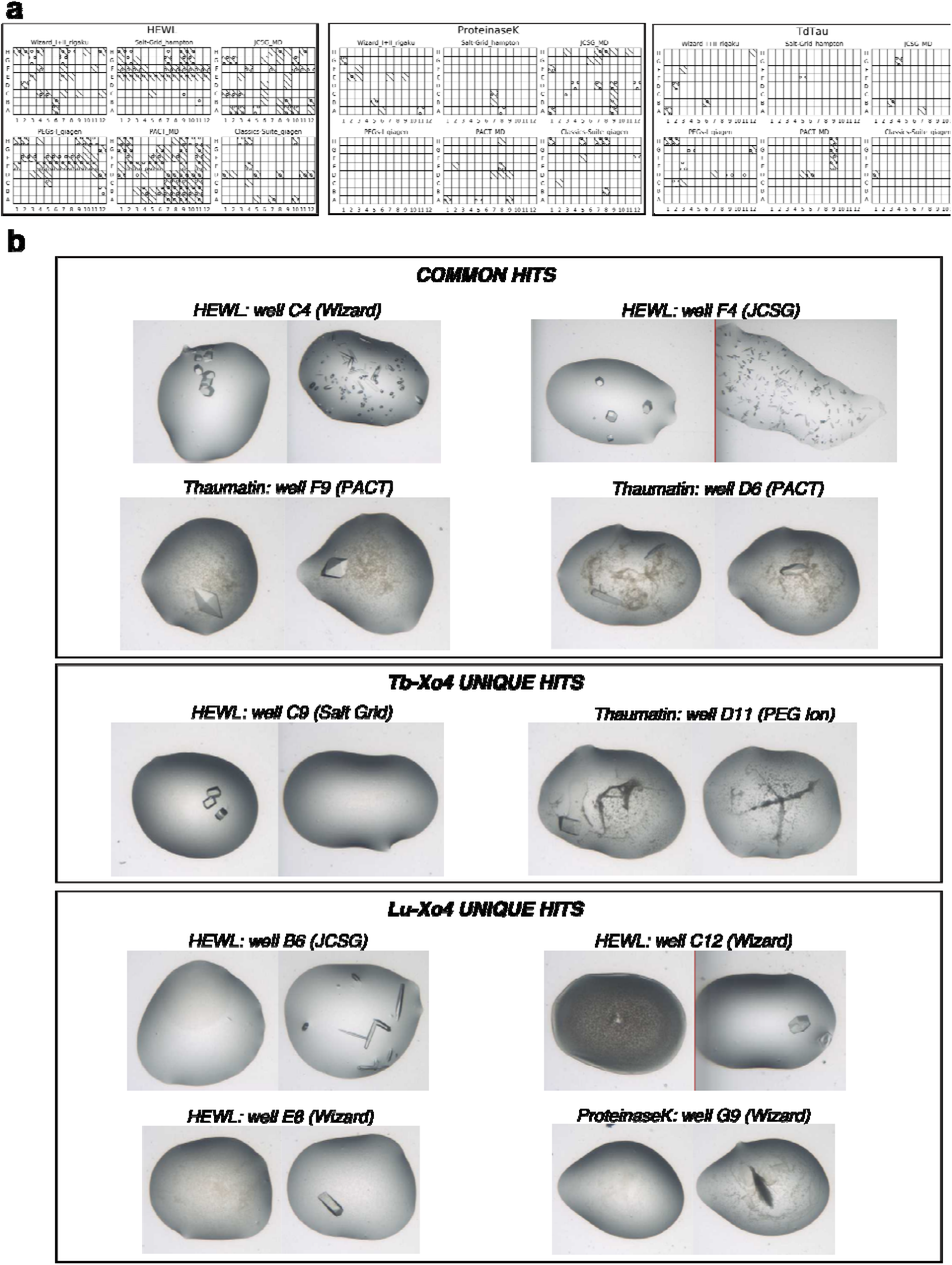
Comparison between the crystallization assays performed with Tb-Xo4 and Lu-Xo4 variants at 10 mM concentration. (a) *By-plate* representations where hits with Tb-Xo4 are depicted by a circle while hits obtained in the presence of Lu-Xo4 are represented with a backslash. (b) Examples of crystallization drops observed in common crystallization conditions (left drops: Tb-Xo4; right drops: Lu-Xo4), in conditions where only Tb-Xo4 leads to crystals and in conditions where crystals only appear in the presence of Lu-Xo4.

### Subset-of-interest representation and exploitation

We also proposed a second way to represent results of crystallization assays. It consists in breaking the *by-plate* level through the definition of a new space called *subset-of-interest* (*SOI*), similarly to the notion of region-of-interest used in image processing for example. In this representation (Figure 4), each crystallization condition leading to a hit is represented by a square. All hit crystallization conditions, for both native and crystallophore assays, are sort out and shared in three categories (−1, 0, or +1) depending of the effect of the crystallophore: (0) the protein crystallizes with and without the crystallophore (it does not affect crystallization), (−1) the protein crystallizes only in absence of the additive (Tb-Xo4 has a negative effect on the crystallization), and (+1) the protein crystallizes only in the presence of the complex (Tb-Xo4 has a positive effect). At this stage, the *SOI* representation could be viewed as an improvement of the histogram representation but it provides a more powerful way to analyze the data as illustrated hereafter.

**Figure 4:**
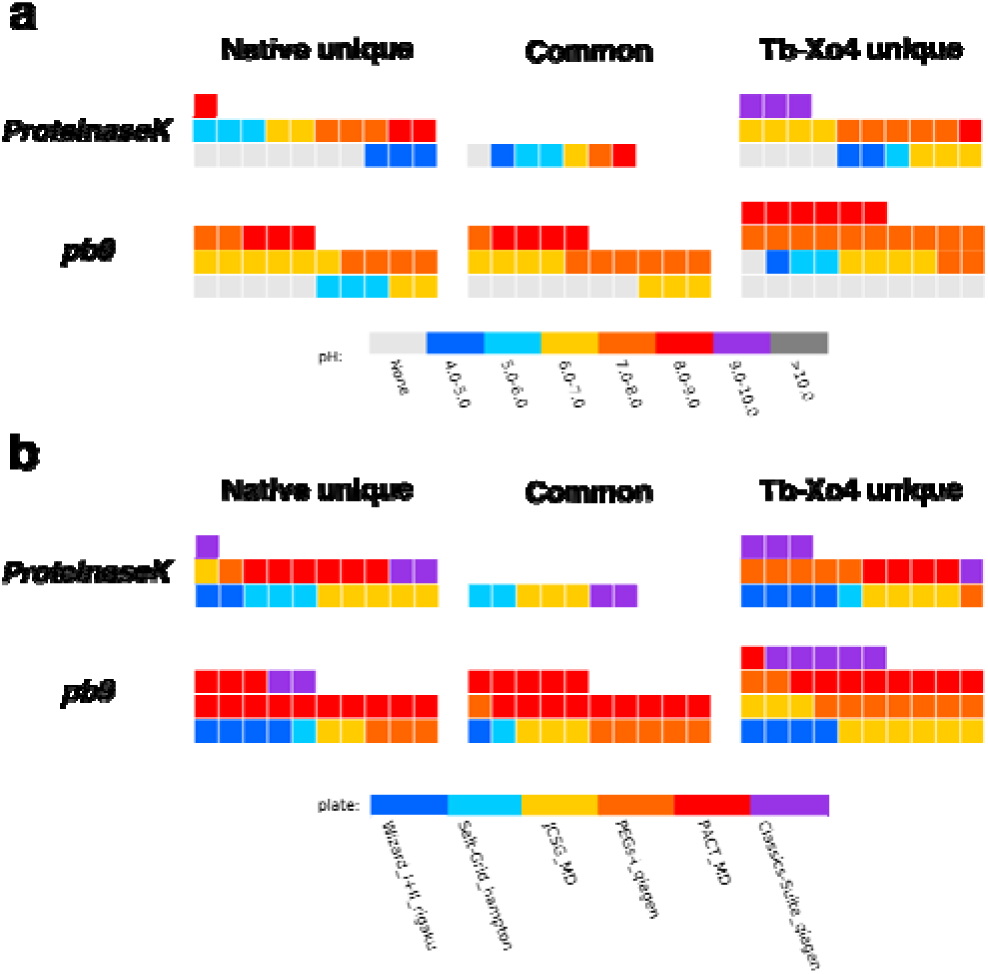
*SOI* representation of the analysis of the influence of (a) pH and (b) crystallization kit on the crystallization of ProteinaseK and pb9 in the absence/presence of 10 mM Tb-Xo4. Each square represents a crystallization hit. The three area of the *SOI* correspond to the hits observed for the protein in the absence/presence of Tb-Xo4 (−1/+1 area, respectively) and observed in both cases (0 area). The reported pH value corresponds to the value of the crystallization conditions available in the description of the commercial kits. *None* indicates that no value is provided in the description.

The presence of a specific parameter can be indicated to highlight possible synergetic or antagonist effects with the additive. Graphically, this can be viewed as a layer on the considered representation (Figure 4) and can be used both in *by-plate* or *SOI* representations. Examples of parameter to be evaluated includes crystallization kit, pH or any crystallization component. In the SOI representation, a given parameter more represented in the -1 area means a detrimental effect with the crystallophore while on the contrary, elements appearing in the +1 will indicate a potential synergetic effect.

Figure 4 illustrates this with two examples of parameters at a single protein level. When pH is considered as parameter, more hits with crystallization solutions with pH above 7 are observed for both proteins in the presence of Tb-Xo4 compared to the protein alone (Figure 4a). In particular, the pH = 9-10 conditions appear only in the +1 area in the case of ProteinaseK clearly indicating a more favorable nucleating condition for the protein in the presence of Tb-Xo4 in such basic medium. We also look at the potential synergy between the crystallophore and a given commercial crystallization kit (Figure 4b). For ProteinaseK when Tb-Xo4 is added, we observed a noticeable increase of the number of hits related to conditions present in the PEGs-Ion kit from Qiagen. A similar synergy with the crystallophore is present for this kit in the case of pb9 as well as an increase of the hits associated with the JCSG kit (Molecular Dimensions). Such analysis are obviously protein-dependent and may be of interest for any future user of crystallophore to determine the best physico-chemical conditions for their protein target to be crystallized. This kind of analysis can also be applied to any other nucleating agents.

However, as part of our ambition to understand the nucleating properties of the crystallophore a larger data set should be considered to ensure meaningful results. The *SOI* analysis shows its full interest when comparative crystallization results of the 6 proteins are taken into account as illustrated in Figure 5. We again considered pH and crystallization kits as parameters.

**Figure 5:**
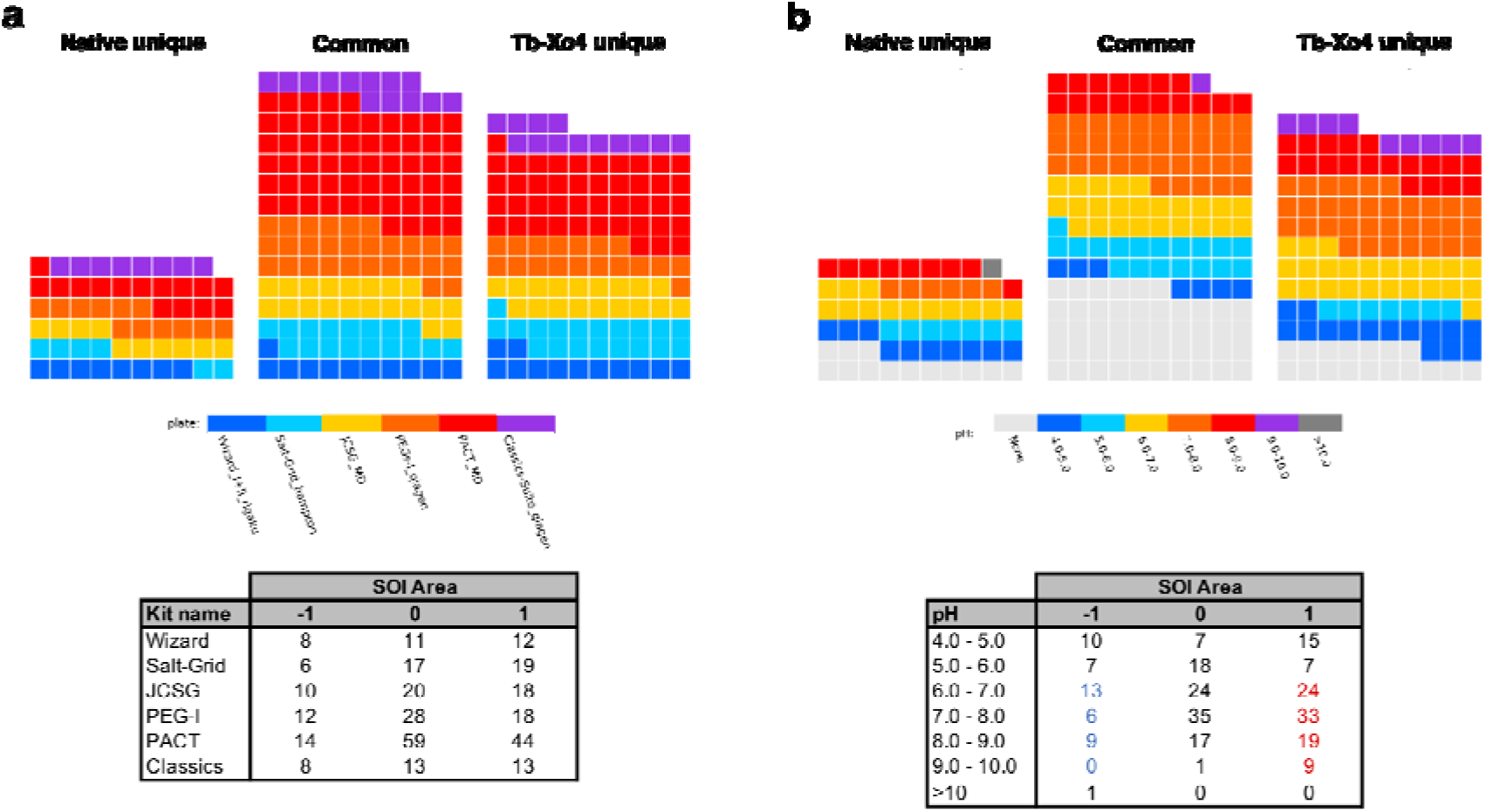
*SOI* representation and analysis (Tables) of the crystallization assays of the 6 considered proteins relative to (a) commercial crystallization kit and (b) pH. Each squar represents a crystallization hit. The three areas of the *SOI* correspond to the hits observed for the protein in the absence/presence of Tb-Xo4 (−1/+1 area, respectively) and observed in both case (0 area). The reported pH value corresponds to the value of the crystallization conditions available in the description of the commercial kits. *None* indicates that no value is provided in the description.

Compared to the conclusion drawn in the case of ProteinaseK and pb9, the analysis performed on the 6 proteins shows a different result on a potential synergy between the crystallophore and a given crystallization kit. We observed a clear increase of the hit number with both Salt-Grid and PACT kit with 3 times more hits when Tb-Xo4 is present (19 and 44 crystallization hits, respectively) compared to in the absence of the crystallophore (Figure 5a). With the basic aim to limit the number of conditions to be evaluated, the analysis performed on the potential synergy between the crystallophore and commercial kits (Figure 5a) shows that the PACT crystallization kit ^36^ maybe a good starting point. Our foreseen studies will obviously focus on the influence of each crystallization components such buffer composition, precipitant nature (PEGs, salts…) or the presence of anions or cations in the crystallization media. Indeed thanks to crystal structure analysis of Tb-Xo4 interactions at the protein surface,^28^ we have observed that components of crystallization solutions, such as sulfate, iodide or magnesium, may directly be involved in the interaction.

The pH tendency observed with ProteinaseK and pb9 proteins is confirmed. More hits (red values in the Table, Figure 5b) are obtained in the presence of Tb-Xo4 when the pH of the crystallization solution is in the range 6-10 compared to native-only hits (blue values in the Table, Figure 5b). The design of Xo4 was done in order to target aspartate and glutamate residues as Tb-Xo4 is a cationic complex. This was later confirmed through crystal structure analysis of Xo4 binding showing a direct coordination of the lanthanide ion by acidic residues,^28^ even though other binding modes were described. Our preliminary observation on the effect of pH are in line with the supposed importance of Xo4 direct coordination. Indeed we observed that the crystallophore favors crystal appearance in crystallization conditions with a pH ranging between 6 and 10. Such pH values ensure that the carboxylate moiety of both aspartate and glutamate are fully deprotonated to ensure a direct interaction with the lanthanide ion(III). This illustrates the added value of setting-up such database. To better take into account the effect of pH and because a non-negligible portion of the commercially available crystallization solutions are provided without any associated pH information, one should get the pH value of each solution by direct measurement or through the use of a dye-based assay as implemented at the CSIRO Collaborative Crystallization Centre (C3).^37^ It would also be of interest to look at correlation between crystallophore nucleating properties and the characteristics of considered proteins such their isoelectric point, the proportion of acidic residues or the ratio between acidic/basic amino acids. For that we plan to introduce a fourth table within the database structure that will contain protein information as depicted in Table 1.

## CONCLUSION

In this article, we present the structure of a database aimed at facilitating the understanding of the unique nucleating properties of a class of lanthanide complexes, the crystallophore. The database exploitation is facilitated by associated tools such as the different representations that have been exemplified. In particular, the concept of *subset-of-interest* has been introduced to reveal potential antagonistic/synergistic effects between Xo4 and other physico-chemical parameters such as pH. The overall approach may be of interest for any studies working on solutions dedicated to improve the nucleating step in protein crystallization.

Thanks to the object-oriented nature of the Python language, as well as the huge amount of available programming libraries, the future development of the database is straight-forward, whether into a stand-alone package or a web-based tool. Finally, the pertinence of such statistical analysis requires that the database is fed with enough occurrences and we invite any users of the crystallophore to share with us the results of their comparative crystallization assays.

## Author Contributions

The manuscript was written through contributions of all authors. All authors have given approval to the final version of the manuscript.

## Notes

OM, FR, EG and SE declare a potential conflict of interest since they are co-founders of the Polyvalan company that commercializes the crystallophore.

## ACKNOWLEDGMENT

IBS acknowledges integration into the Interdisciplinary Research Institute of Grenoble (IRIG, CEA). This work used the platforms of the Grenoble Instruct-ERIC center (ISBG; UMS 3518 CNRS-CEA-UGA-EMBL) within the Grenoble Partnership for Structural Biology (PSB), supported by FRISBI (ANR-10-INBS-05-02) and GRAL, financed within the University Grenoble Alpes graduate school (Ecoles Universitaires de Recherche) CBH-EUR-GS (ANR-17- EURE-0003). Authors also thank the pôle de compétitivité Lyon-Biopôle. Authors are grateful to Christèle Maury for fruitful discussion in the elaboration of the SOI analysis.

## ABBREVIATIONS

Xo4: crystallophore;
SOI: subset-of-interest;
HEWL: Hen egg white lysozyme;
GRHPR: Glyoxylate Hydroxypyruvate Reductase.

